# Sex-Specific Role of Myostatin Signaling in Neonatal Muscle Growth, Denervation Atrophy, and Neuromuscular Contractures

**DOI:** 10.1101/2022.06.17.496582

**Authors:** Marianne E Emmert, Parul Aggarwal, Kritton Shay-Winkler, Se-Jin Lee, Qingnian Goh, Roger Cornwall

**Affiliations:** Department of Medical Sciences, University of Cincinnati College of Medicine, Cincinnati, OH, USA; Division of Orthopaedic Surgery, Cincinnati Children’s Hospital Medical Center, Cincinnati, OH, USA; The Jackson Laboratory, Farmington, CT, USA; Department of Genetics and Genome Sciences, University of Connecticut School of Medicine, Farmington, CT, USA; Department of Orthopaedic Surgery, University of Cincinnati College of Medicine, Cincinnati, OH, USA; Division of Developmental Biology, Cincinnati Children’s Hospital Medical Center, Cincinnati, OH, USA; Department of Pediatrics, University of Cincinnati College of Medicine, Cincinnati, OH, USA

**Author notes:** To whom correspondence should be addressed: Qingnian Goh, PhD, Cincinnati Children’s Hospital Medical Center 3333 Burnet Ave, MLC 2017, Cincinnati, OH 45229; Or to: Roger Cornwall, MD, Cincinnati Children’s Hospital Medical Center 3333 Burnet Ave, MLC 2017, Cincinnati, OH 45229.

**Keywords:** Sex dimorphisms, myostatin, neonatal brachial plexus injury, denervation, neuromuscular contractures, muscle growth, muscle atrophy, muscle length, proteostasis

## Abstract

Neonatal brachial plexus injury (NBPI) causes disabling and incurable muscle contractures that result from impaired longitudinal growth of denervated muscles. This deficit in muscle growth is driven by increased proteasome-mediated protein degradation, suggesting a dysregulation of muscle proteostasis. The myostatin (MSTN) pathway, a prominent muscle-specific regulator of proteostasis, is a putative signaling mechanism by which neonatal denervation could impair longitudinal muscle growth, and thus a potential target to prevent NBPI-induced contractures. Through a mouse model of NBPI, our present study revealed that pharmacologic inhibition of MSTN signaling induces hypertrophy, restores longitudinal growth, and prevents contractures in denervated muscles of female but not male mice, despite inducing hypertrophy of normally innervated muscles in both sexes. Additionally, the MSTN-dependent impairment of longitudinal muscle growth after NBPI in female mice is associated with perturbation of 20S proteasome activity, but not through alterations in canonical MSTN signaling pathways. These findings reveal a sex dimorphism in the regulation of neonatal longitudinal muscle growth and contractures, thereby providing insights into contracture pathophysiology, identifying a potential muscle-specific therapeutic target for contracture prevention, and underscoring the importance of sex as a biological variable in the pathophysiology of neuromuscular disorders.

## Introduction

Injury to the brachial plexus at birth (Neonatal Brachial Plexus Injury – NBPI) is the most common cause of upper limb paralysis in children,^1^ occurring in approximately 1.5 of every 1,000 live births. This initial nerve injury leads to permanent neurologic deficits in 20-40% of affected children,^2,3^ and results in the secondary formation of disabling and incurable muscle contractures, or joint stiffness. Contractures severely impede range of motion and functional use of the involved limbs, ultimately resulting in skeletal deformity that further worsens limb dysfunction.^4^ However, current strategies are insufficient in restoring muscle function and joint range of motion once contractures have developed, and may even worsen function by further weakening abnormal muscles.^5–7^ To develop effective strategies for treating contractures, it is therefore important to establish greater insights into the pathophysiology of contracture formation.

Using a mouse model of NBPI, we previously discovered that contractures result from impaired longitudinal muscle growth, as characterized by the overstretch/elongation of sarcomeres in denervated muscles.^8–11^ Such deficits in muscle length are caused by aberrant levels of muscle protein degradation associated with increased catalytic activity of the 20S proteasome.^12^ These results posit a critical mechanistic role for proteasome-mediated dysregulation of muscle proteostasis in driving the impairment of longitudinal muscle growth that ultimately causes contractures. As proof of concept, we showed that inhibition of the proteasome with an FDA-approved proteasome inhibitor, bortezomib, restores sarcomere length and reduces contracture formation.^12^ Our collective findings therefore establish that the biomechanical contracture phenotype can be attributed to a biological defect, and that contractures can be corrected by targeting their causative mechanism. However, despite the effectiveness of proteasome inhibition for preventing contractures, this pharmacologic strategy cannot be easily translated to children. We recently reported that bortezomib treatment is required throughout postnatal development to prevent contractures at skeletal maturity, and that its efficacy is diminished beyond the neonatal period.^13^ This need for chronic bortezomib treatment is problematic. Prolonged administration of proteasome inhibitors results in potential cumulative toxicity as these drugs nonspecifically block degradation and cause tissue damage to many organs, such as the brain,^12^ and even impede musculoskeletal development.^13^ Due to these concerns, our lab seeks to identify safer strategies to prevent denervation-induced contractures by targeting skeletal muscle-specific regulators of proteostasis to reduce the elevated protein degradation responsible for contractures.

One such signaling mechanism specific to skeletal muscle is the myostatin (MSTN) pathway. MSTN, also known as growth differentiation factor-8 (GDF8), is a myokine and a member of the transforming growth factor-β (TGF-β) superfamily.^14^ In normally innervated skeletal muscles, ligand binding of MSTN to Activin A through its Type 2 receptors (ACVR and ACVR2B) activates the downstream Smad proteins 2 & 3.^15,16^ These signaling events regulate muscle homeostasis by increasing degradation and decreasing Akt/mTOR-mediated synthesis, ultimately attenuating muscle size and limiting aberrant growth.^14,17,18^ MSTN thereby functions as a negative regulator of muscle mass and protein balance. Given its prominent role in muscle proteostasis, it is possible that MSTN is a signaling pathway by which denervation impairs muscle growth. Specifically, we speculate that neonatal denervation induces contractures through MSTN-dependent impairment of postnatal longitudinal muscle growth. If contractures are indeed mediated through MSTN signaling, it could lead to a potential breakthrough in identifying a muscle-specific target for contracture prevention.

Hence, in the current study, we seek to elucidate the role of MSTN in the formation of denervation-induced muscle contractures. Using a soluble decoy receptor (ACVR2B-Fc) to inhibit ligand binding of MSTN to Activin A,^19–22^ we specifically investigated whether pharmacologic inhibition of MSTN signaling preserves longitudinal muscle growth and prevents contractures after NBPI. Our collective results establish several novel insights in skeletal muscle biology and contracture pathophysiology. First, we show that pharmacologic MSTN inhibition is efficacious in augmenting neonatal growth of normally innervated skeletal muscles in both female and male mice. Second, MSTN inhibition effectively restores sarcomere length and reduces contracture severity exclusively in neonatally denervated muscles of female mice, suggesting a sex-dependent role for the MSTN pathway in modulating longitudinal muscle growth and contracture formation. Curiously, MSTN inhibition improves muscle proteostasis in denervated female muscles by circumventing known downstream signaling pathways to directly target the 20S proteasome. These discoveries establish a critical link between denervation and impaired longitudinal muscle growth, which helps guide translation of safer strategies for preventing contractures. Furthermore, they highlight the importance of sex as a biological variable in disease pathology and treatment strategies, as well as in future studies dissecting the molecular regulation of longitudinal muscle growth.

## Results

### MSTN inhibition enhances neonatal skeletal muscle growth

While pharmacologic inhibition of MSTN signaling with the soluble ACVR2B-Fc decoy receptor has been shown to induce robust muscle growth in adult mice, it has not been validated in neonatal mice. To limit potential toxicity during the neonatal period, we standardized the frequency of ACVR2B-Fc treatment to weekly injections.^19,20^ We began by verifying the effects of this dosage on neonatal skeletal muscle growth (**Figure 1A**). To further account for sexual dimorphisms in muscle size during postnatal development,^23^ we analyzed the effect of MSTN inhibition on developmental growth according to individual sexes. Our findings revealed that inhibition of the MSTN pathway resulted in robust growth of normally innervated forelimb muscles in both neonatal female and male mice. Specifically, MSTN inhibition enhanced cross-sectional and volumetric growth of the brachialis muscles (**Figures 1B-D**), as well as increased overall muscle weight and elevated total protein levels in biceps muscles (**Figures 1F-G**). Despite this, the increases in brachialis volume (61% vs. 35%), biceps weight (49% vs. 35%), and biceps protein content (47% vs. 28%) was greater in females than males when compared to respective controls (**Figures 1E, 1H**). These results indicate that while neonatal inhibition of MSTN signaling effectively enhances skeletal muscle growth in both sexes, it facilitates greater protein anabolism and growth in female muscles. Curiously, ACVR2B-Fc treatment did not yield additional gains in body weight but instead attenuated humerus length and heart size in male mice, whereas female mice had smaller spleens after treatment (**Figure 1 – supplemental figures 1A-D**). Hence, while pharmacologic MSTN inhibition is associated with several off-target effects, these alterations manifest more prevalently in male tissues.

**Figure 1:**
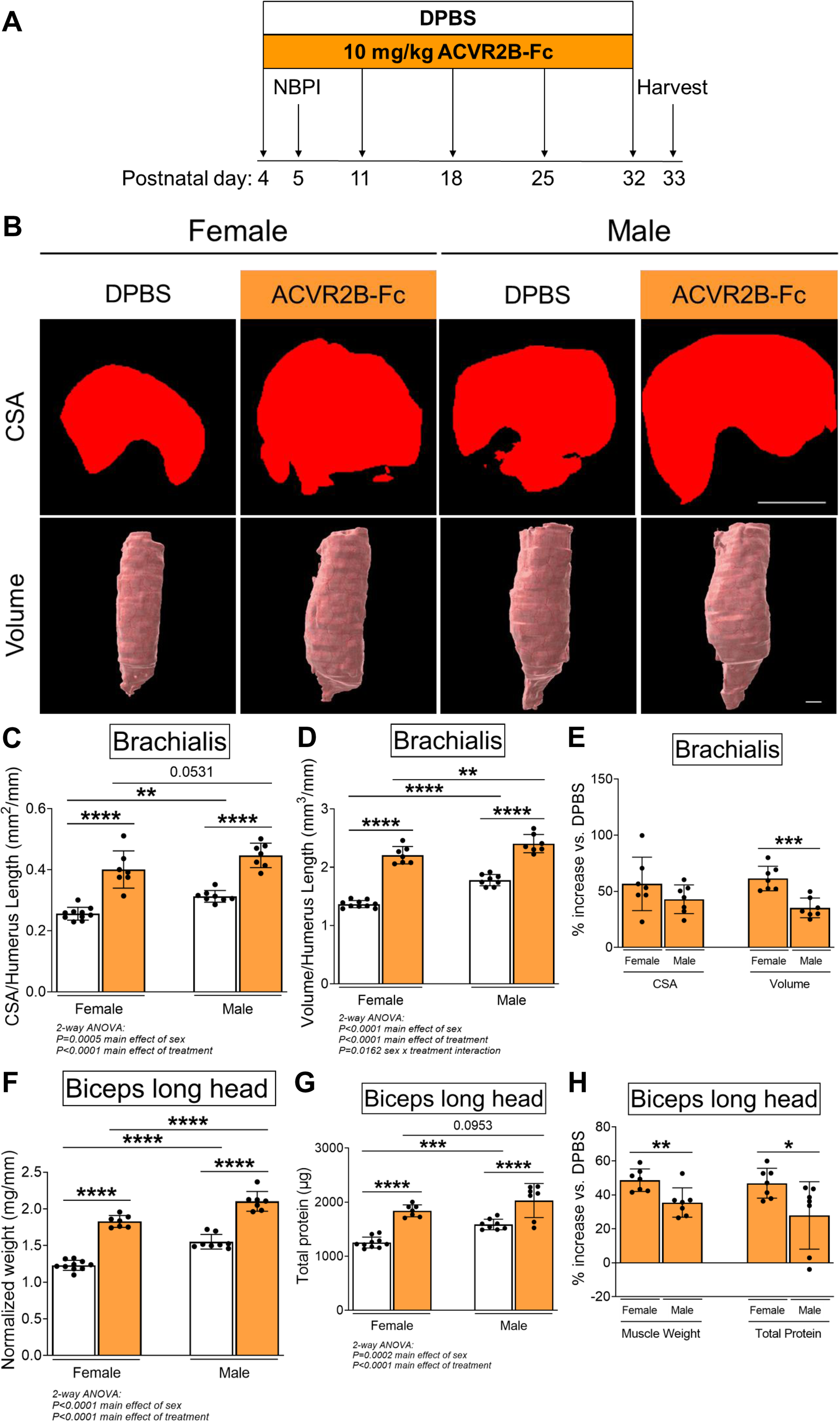
Pharmacologic inhibition of MSTN enhances neonatal skeletal muscle growth. (A) Schematic depiction of ACVR2B-Fc treatment to inhibit MSTN in neonatal mice prior to and after unilateral brachial plexus injury (NBPI) at P5. Representative micro-CT images in (B) transverse and 3-dimensional views revealed increased growth of non-denervated brachialis muscles in both female and male mice following 4 weeks of pharmacologic MSTN inhibition with ACVR2B-Fc. Quantitative analyses of (C) cross-sectional area and (D) whole muscle volume in control brachialis muscles confirmed that neonatal MSTN inhibition enhances skeletal muscle growth. Neonatal MSTN inhibition further enhances (F) muscle weight and (G) total protein content of non-denervated biceps muscles in both sexes. Despite this, (E), (H) the increases in muscle volume, muscle weight, and protein levels were larger in females than males when compared to their respective DPBS controls. Data are presented as mean ± SD, n = 7-10 independent mice. Statistical analyses: (C), (D), (F), (G) 2-way ANOVA for sex and treatment with a Bonferroni correction for multiple comparisons, (E), (H) unpaired two-tailed Student’s t-tests. **\***P<0.05, **\*\***P<0.01, **\*\*\***P<0.001, **\*\*\*\***P<0.0001. Scale bar: 1000 µm.

### MSTN signaling modulates denervation atrophy in a sex-specific manner

To elucidate the contribution of the MSTN pathway in modulating contracture formation, we began by investigating the effects of ACVR2B-Fc treatment on neonatally denervated forelimb muscles. In contrast to normally innervated neonatal muscles, pharmacologic blockade of MSTN signaling reduced muscle loss of denervated brachialis and biceps muscles only in female mice (**Figures 2A-E**). In denervated female muscles, MSTN inhibition specifically increased brachialis cross-sectional area and volume by 25-26%, and biceps weight and total protein by 31-50%. However, there were no improvements with MSTN inhibition in denervated male muscles, as ACVR2B-Fc treatment failed to increase brachialis muscle CSA and volume (**Figures 2A-C**), or biceps muscle weight and total protein levels (**Figures 2D-E**) in male mice. This lack of effect of neonatal MSTN inhibition on denervated muscles in male mice was further associated with an attenuation of skeletal growth in the denervated humeri (**Figure 2 – supplemental figure 1A**). Taken together, our results illustrate that while neonatal MSTN inhibition facilitates developmental growth of normally innervated muscles in both males and females, it promotes neonatal growth and reduces atrophy of denervated muscles exclusively in females.

**Figure 2:**
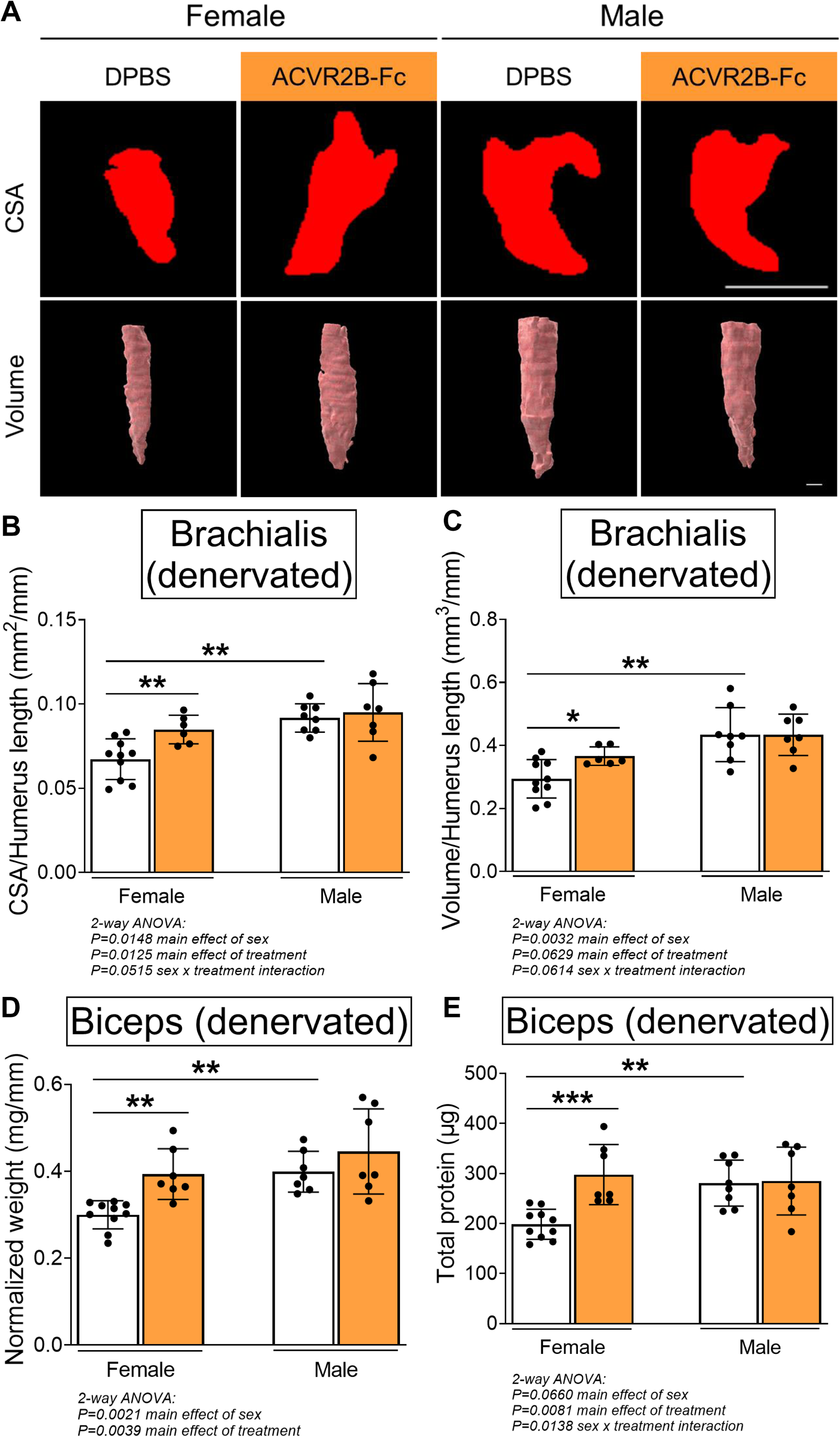
Pharmacologic MSTN inhibition reduces denervation-induced muscle atrophy in neonatal female mice. Representative micro-CT images in (A) transverse and 3-dimensional views revealed larger denervated brachialis muscles in female mice following MSTN inhibition. Quantitative analyses of (B) brachialis cross-sectional area and (C) whole muscle volume confirmed MSTN inhibition reduces denervation-induced atrophy only in female mice. This partial rescue in growth of the denervated brachialis muscles is accompanied by increased (D) muscle weight and (E) total protein content of denervated biceps muscles in female mice only. Data are presented as mean ± SD, n = 7-10 independent mice. Statistical analyses: (B), (C), (D), (E) 2-way ANOVA for sex and treatment with a Bonferroni correction for multiple comparisons. **\***P<0.05, **\*\***P<0.01, **\*\*\***P<0.001. Scale bar: 1000 µm.

### MSTN signaling modulates contracture formation in a sex-specific manner

Due to the differential effects in denervated muscle growth, we speculated that the MSTN signaling pathway may mediate contracture formation in a sex-dependent manner. Indeed, we discovered that ACVR2B-Fc treatment improved passive range of motion in both elbow and shoulder joints only in female mice 4 weeks post-NBPI (**Figures 3A-B, 3D-E**). This sex-specific improvement in joint mobility corresponded to a reduction in both elbow flexion and shoulder rotation contracture severity in female, but not male mice (**Figures 3C, 3F**). These results indicate a critical therapeutic window for MSTN inhibition in preventing contracture formation, and importantly, illuminate a sex-specific role for the MSTN pathway in mediating contracture formation. To gain further insights on this sex dimorphism, we next assessed the role of MSTN signaling on longitudinal growth of denervated muscles. Here, we observed that neonatal MSTN inhibition rescued sarcomere length (reduced sarcomere elongation) in the denervated brachialis muscles of female mice (**Figures 4A-C**), indicating an improvement in functional muscle length.^13^ Conversely, sarcomeres in the denervated brachialis of male mice remained overstretched, an indication of impaired longitudinal muscle growth. Hence, our collective findings establish that contracture formation following neonatal muscle denervation in female mice is driven by MSTN-dependent impairment of longitudinal muscle growth.

**Figure 3:**
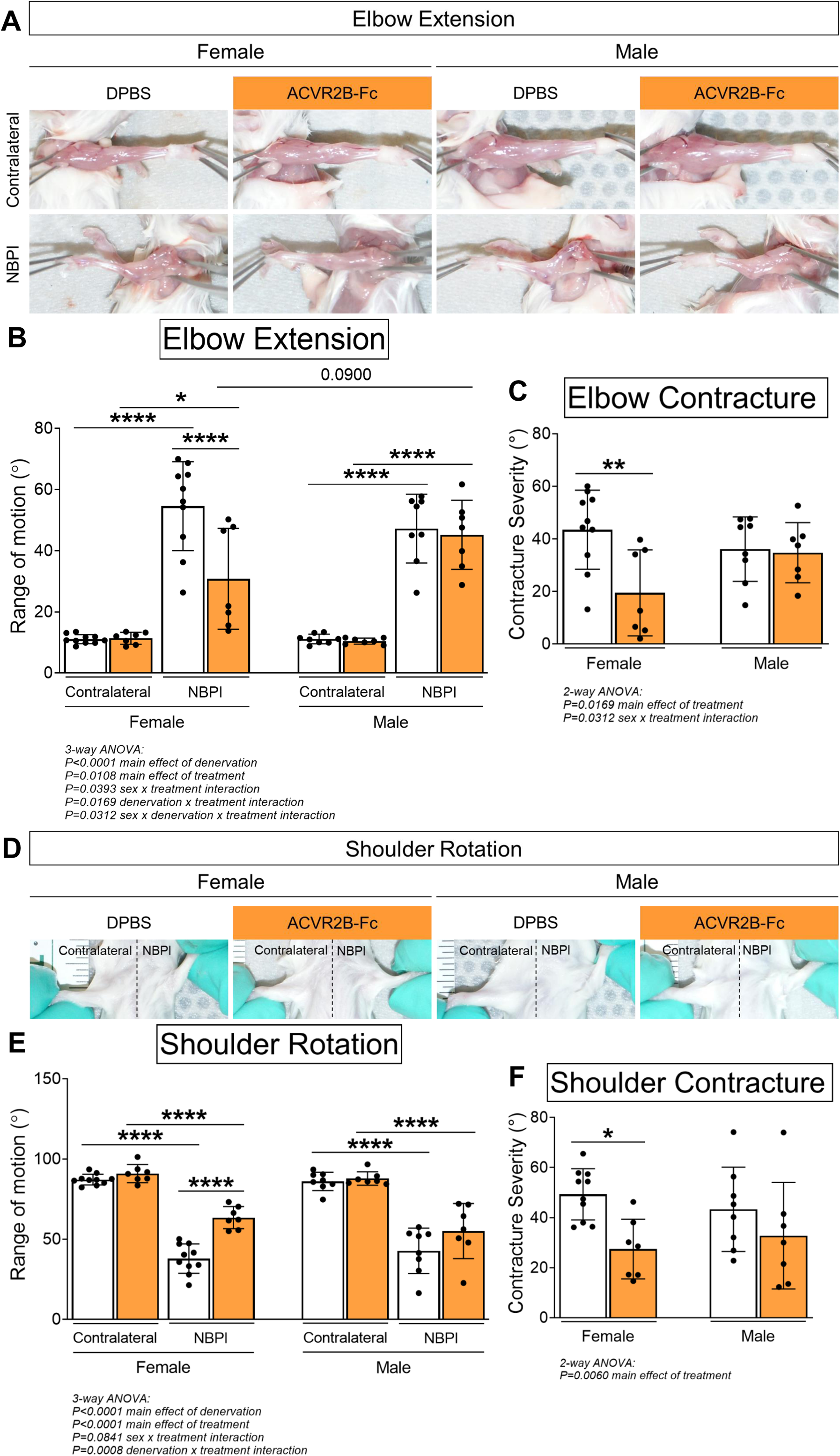
Pharmacologic MSTN inhibition reduces neonatal denervation-induced contractures in neonatal female mice. (A) Representative images of denervated (NBPI) and contralateral forelimbs, and quantitation of both (B) elbow joint range of motion and (C) contracture severity revealed that MSTN inhibition reduces the formation of elbow flexion contractures after 4 weeks of neonatal denervation in female, but not male mice. Similarly, (D) representative images of bilateral forelimbs, and quantitation of both (E) shoulder joint range of motion and (F) contracture severity demonstrated improvements in shoulder rotation only in female mice. In (C) and (F), elbow and shoulder contracture severity is calculated as the difference in passive elbow extension and shoulder rotation, respectively, between the NBPI side and the contralateral side. Data are presented as mean ± SD, n = 7-10 independent mice. Statistical analyses: (B), (E) 3-way ANOVA for sex, treatment, and denervation (repeated measures between forelimbs) with a Bonferroni correction for multiple comparisons, (C), (F) 2-way ANOVA for sex and treatment with a Bonferroni correction for multiple comparisons. **\***P<0.05, **\*\***P<0.01, **\*\*\*\***P<0.0001.

**Figure 4:**
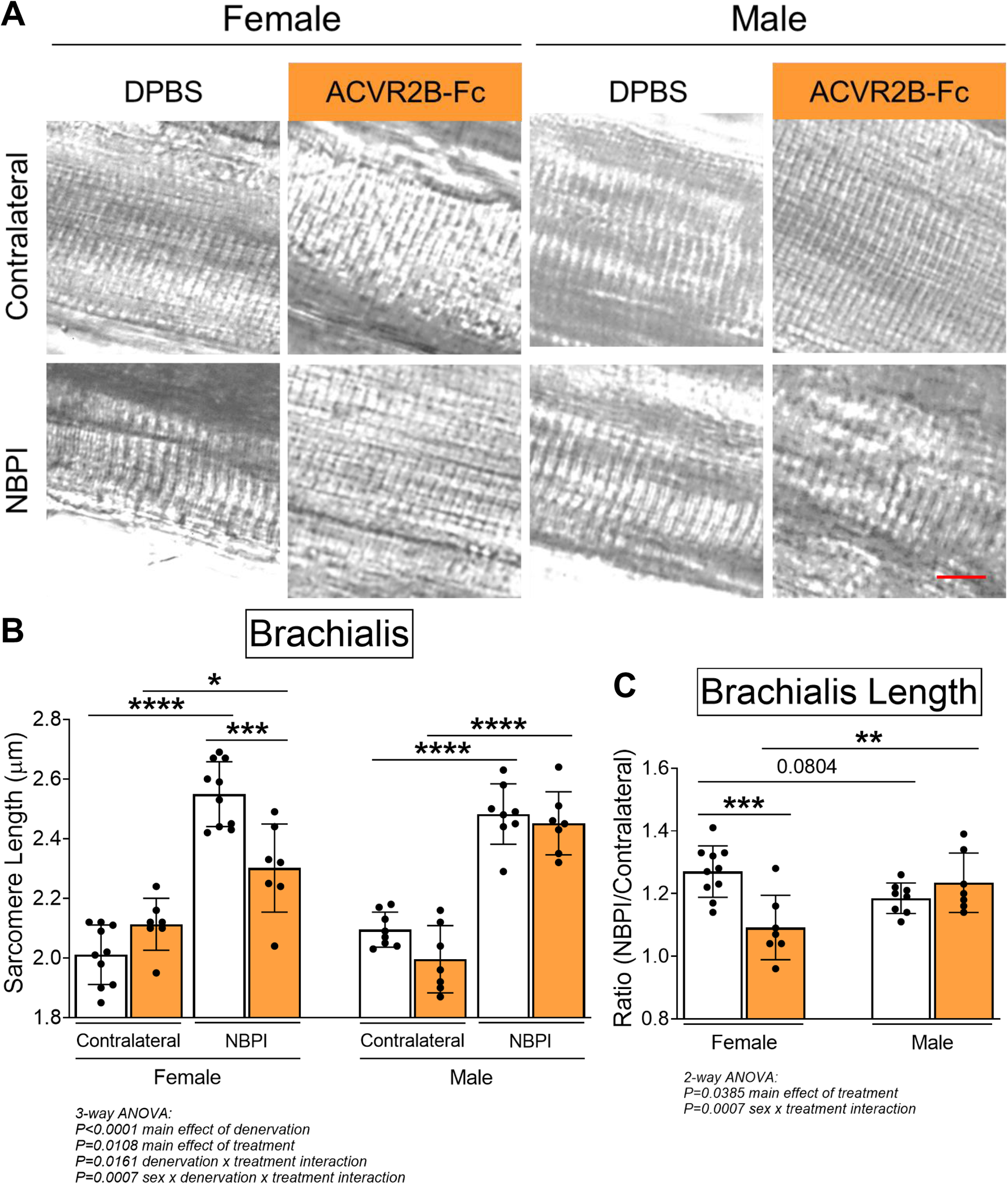
Pharmacologic MSTN inhibition preserves longitudinal muscle growth of denervated muscles in neonatal female mice. (A) Representative DIC images of sarcomeres, and (B) quantitation of sarcomere length showed that MSTN inhibition preserves sarcomere length of denervated muscles after neonatal denervation in female mice, whereas the sarcomeres of denervated muscles in male mice remain overstretched. (C) Sarcomere length on the NBPI side is normalized to the contralateral side to generate a sarcomere length ratio. This normalized sarcomere length between control and NBPI limbs further confirmed that MSTN inhibition improves functional length of denervated muscles only in female mice. Data are presented as mean ± SD, n = 7-10 independent mice. Statistical analyses: (B) 3-way ANOVA for sex, treatment, and denervation (repeated measures between forelimbs) with a Bonferroni correction for multiple comparisons, (C) 2-way ANOVA for sex and treatment with a Bonferroni correction for multiple comparisons. **\***P<0.05, **\*\***P<0.01, **\*\*\***P<0.001, **\*\*\*\***P<0.0001. Scale bar: 10 µm.

### MSTN mediates denervation-induced proteostasis dysregulation in a sex-specific manner

Since our prior work revealed that impaired longitudinal muscle growth following neonatal denervation is driven by increased levels of proteasome-mediated protein degradation,^12^ we subsequently explored the role of MSTN signaling in proteostasis dysregulation to decipher mechanisms governing the observed sex dimorphisms. We first assessed whether MSTN inhibition is able to further increase the elevated levels of protein synthesis in neonatally denervated muscles.^12^ Through western blot analysis of puromycin incorporation,^24,25^ we discovered that while 4 weeks of NBPI drove protein synthesis in denervated biceps muscles, MSTN inhibition does not augment this increase in either sex (**Figures 5A-B**). This finding was corroborated by western blot analyses of the Akt/mTOR hypertrophic signaling pathway,^26–29^ which revealed that perturbations in Akt activity and total protein expression in denervated triceps muscles were not further impacted by ACVR2B-Fc treatment at 4 weeks post-NBPI (**Figures 5C-F**). These collective findings therefore indicate that MSTN-mediated proteostasis dysregulation in denervated female muscles occurs independent of Akt/mTOR-mediated protein synthesis.

**Figure 5:**
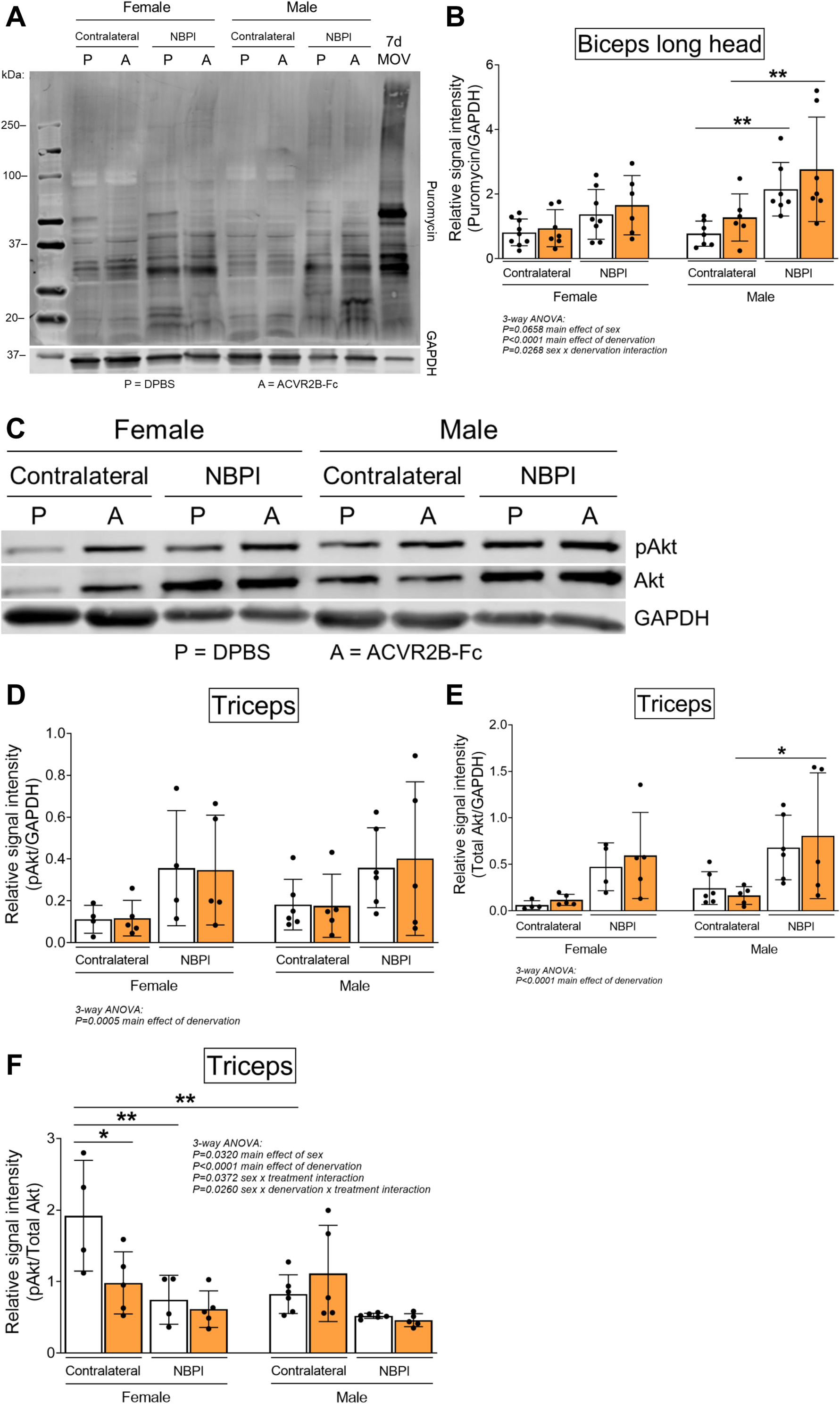
Sex-specific differences in MSTN-dependent contracture formation is not mediated through protein synthesis. (A) Representative western blots of puromycin incorporation and (B) quantitative analysis of optical densities revealed MSTN inhibition does not further alter the denervation-induced increases in whole muscle protein synthesis in biceps muscles of both female and male mice. n = 7-10 independent mice. (C) Representative western blots and quantitative analyses of (D) pAkt (Ser473) and (E) total Akt similarly showed that ACVR2B-Fc treatment does not lead to additional increases in activity and translation of Akt after denervation in both sexes. (F) Quantification of the western signal for pAkt normalized to total protein levels further indicated that MSTN inhibition does not alter Akt/mTOR signaling in neonatally denervated muscles. n = 4-6 independent mice. Statistical analyses: (B), (D), (E), (F) 3-way ANOVA for sex, treatment, and denervation (repeated measures between forelimbs) with a Bonferroni correction for multiple comparisons. **\***P<0.05, **\*\***P<0.01. 7d MOV = Adult mouse plantaris muscle that had been subjected to 7 days of mechanical overload.

Having ruled out synthesis as an underlying mechanism for the observed sex dimorphisms, we next investigated the role of degradation, the other side of proteostasis. While total levels of K48-linked polyubiquitin were not different between normally innervated and denervated biceps muscles, we observed increased polyubiquitination of proteins greater than 40 kDa (**Figures 6A-D**), similar to what we have reported before.^12^ However, MSTN inhibition did not attenuate the denervation-induced increases in proteins tagged for degradation at this molecular weight range in either sex. Since MSTN is known to activate the ubiquitin-proteasome pathway by upregulating the expression of upstream ubiquitin ligases,^30,31^ it was surprising that its inhibition did not reduce ubiquitination levels. Instead, ACVR2B-Fc effectively reduced the catalytic activity of the 20S proteasome β5 subunit in denervated triceps muscles only in female mice (**Figures 6E-F**). As K48-linked ubiquitin chains are the most prevalent signal that mark protein substrates for proteasome degradation,^32^ these findings suggest that MSTN regulates sex-specific proteostasis dysregulation by circumventing polyubiquitination and directly targeting the proteasome. To further decipher signaling mechanisms governing the sex differences in MSTN-mediated protein degradation, we characterized the Smad2/3 pathway in triceps muscles.^28,29,33,34^ Here, we observed sex-independent increases in Smad3 phosphorylation and translation with neonatal denervation, which were not further altered with MSTN inhibition (**Figures 7A-D**). Our collective findings thus posit a sex-specific role for MSTN in muscle proteostasis dysregulation after neonatal denervation through non-canonical signaling pathways.

**Figure 6:**
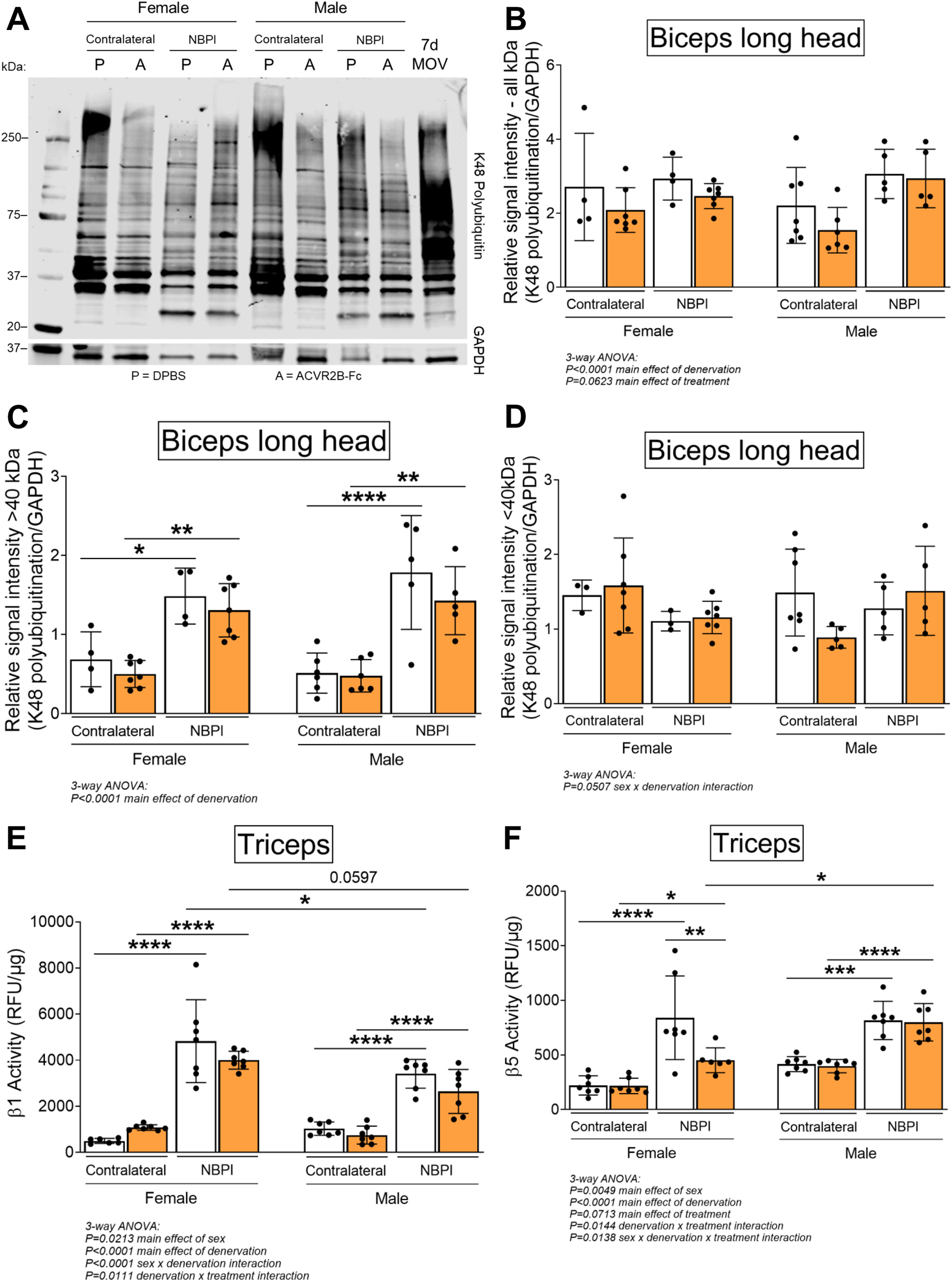
Sex-specific differences in MSTN-dependent contracture formation is mediated through proteasome activity. (A) Representative western blots of K48-linked polyubiquitin and (B) quantitative analysis of optical densities showed similar levels of ubiquitination in control and denervated biceps muscles of neonatal female and male mice. Despite this, (C) in-depth analyses of optical densities discovered that denervation increases ubiquitination levels of higher molecular weight proteins >40 kDa, (D) but not lower molecular proteins <40 kDa in both sexes. Importantly, MSTN inhibition does not alter levels of K48 polyubiquitination across the different molecular weights after denervation. n = 4-7 independent mice. (E), (F) Assessment of proteasome activity in triceps muscles revealed that MSTN inhibition blunts the denervation-induced increase in β5 but not β1 constitutive proteasome activity solely in female mice. n = 7 independent mice. Data are presented as mean ± SD. Statistical analyses: (B), (C), (D), (E), (F) 3-way ANOVA for sex, treatment, and denervation (repeated measures between forelimbs) with a Bonferroni correction for multiple comparisons. **\***P<0.05, **\*\***P<0.01, **\*\*\***P<0.001, **\*\*\*\***P<0.0001. 7d MOV = Adult mouse plantaris muscle that had been subjected to 7 days of mechanical overload.

**Figure 7:**
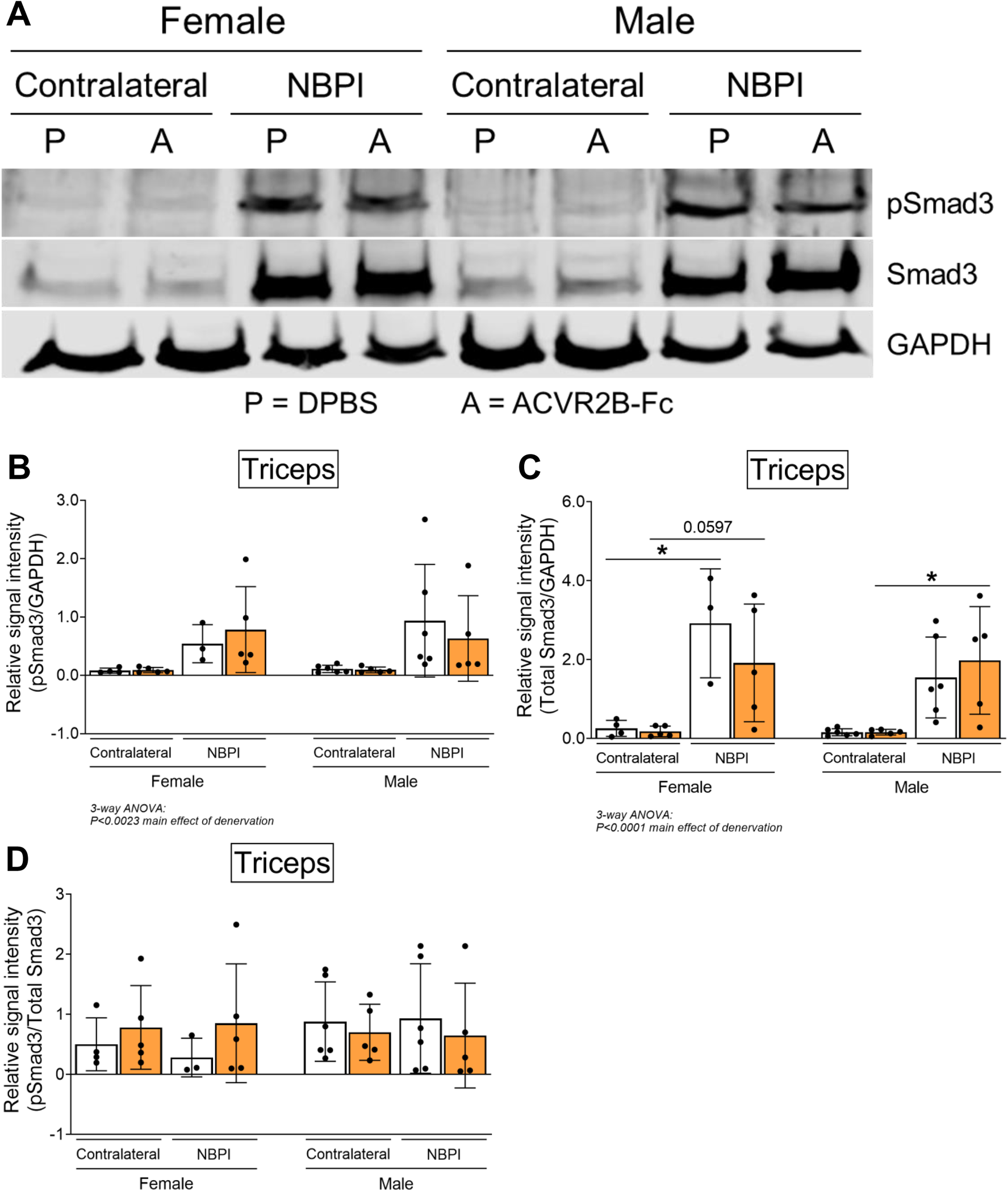
Sex-specific differences in MSTN-mediated proteostasis dysregulation occurs independent of Smad2/3 signaling. (A) Representative western blots and quantitative analyses of (B) pSmad3 and (C) total Smad3 revealed that ACVR2B-Fc treatment does not blunt the denervation-induced increase in activity and translation of Smad3 in triceps muscles of both sexes. (D) Quantification of the western signal for pSmad3 normalized to total protein levels further indicated that MSTN inhibition does not alter Smad2/3 signaling in neonatally denervated muscles. n = 3-6 independent mice. Data are presented as mean ± SD. Statistical analyses: (B), (C), (D) 3-way ANOVA for sex, treatment, and denervation (repeated measures between forelimbs) with a Bonferroni correction for multiple comparisons. **\***P<0.05.

### Less pronounced sex differences with proteasome inhibition

Having observed a role for sex in MSTN-dependent contracture formation, we revisited our prior findings of long-term proteasome inhibition in preventing contractures.^13^ Here, we reanalyzed our data from 4, 8, and 12w of continuous bortezomib treatment (**Figure 8A**) according to sex. In contrast with MSTN inhibition, we observed greater and more consistent reductions in elbow and shoulder contracture severity across the different time points in male mice with bortezomib treatment (**Figures 8B-C**). Sex differences in sarcomere length were less apparent, as bortezomib improved longitudinal muscle growth in female and male mice, except at 8w of treatment (**Figure 8D**). Lastly, while bortezomib did not alter β5 subunit activity in either sex, it attenuated β1 subunit activity at 8 and 12w only in male mice (**Figures 8E-F**). Our prior reports of reduced β1 activity with bortezomib treatment at 4w were observed only in female mice, as proteasome activity was not measured in male mice (data not shown).^13^ Overall, while sex differences do exist with neonatal and long-term bortezomib treatment, these differences are more subtle than sex dimorphisms observed with MSTN inhibition. Regardless, they further highlight that sex mediates neonatally-induced contractures through divergent pathways.

**Figure 8:**
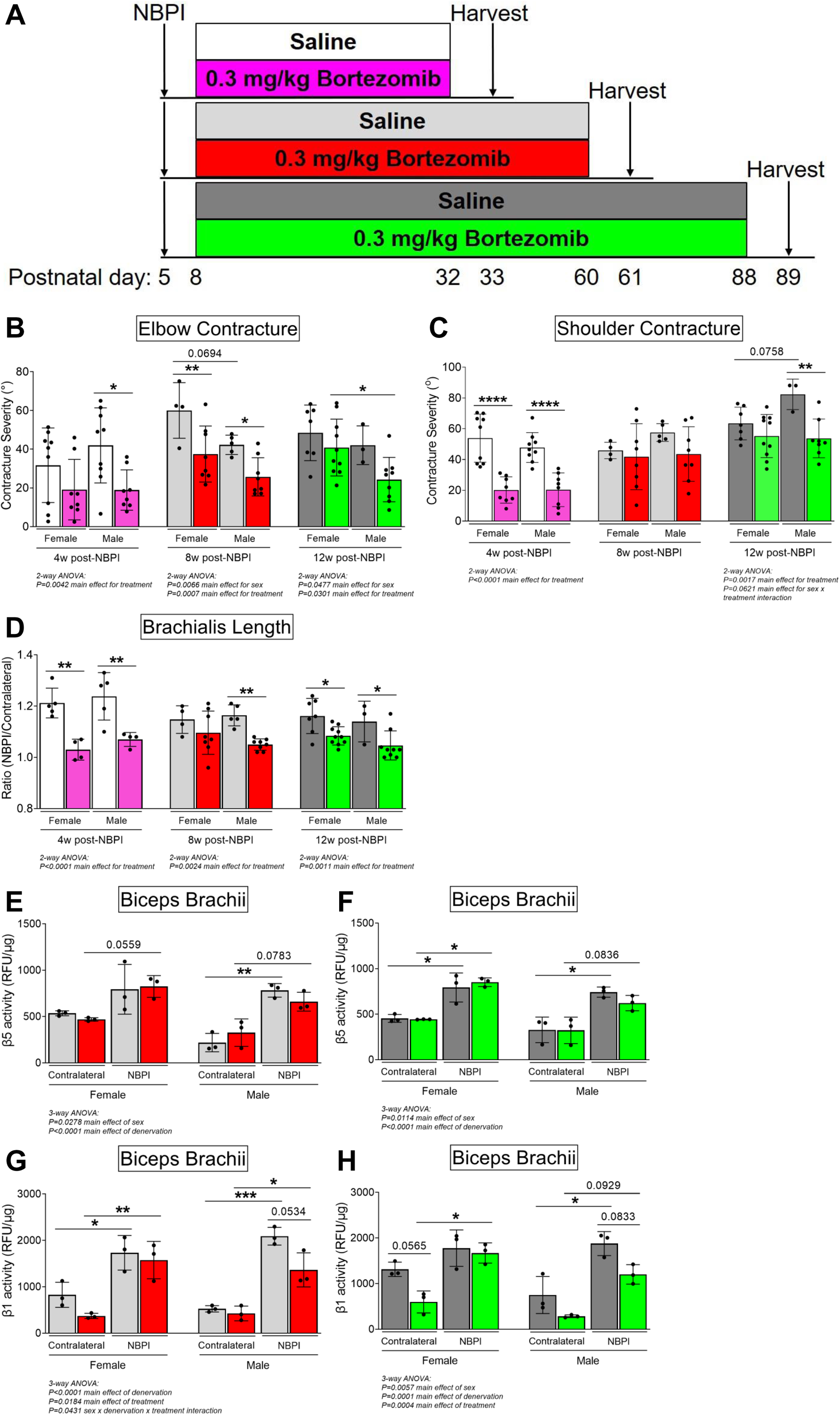
Proteasome inhibition with bortezomib prevents contractures and restores proteostasis preferentially in denervated muscles of neonatal male mice. (A) Schematic depiction of bortezomib treatment and assessment of outcomes at 4, 8, or 12 weeks post-NBPI. (B), (C) Continuous bortezomib treatment preferentially prevents elbow and shoulder contractures in male mice throughout postnatal development and into skeletal maturity (n = 3-10 independent mice). (D) Despite this, long-term bortezomib does not confer sex-specific improvements in longitudinal muscle growth, as sarcomere length was preserved in denervated brachialis muscles of both female and male mice (n = 3-10 independent mice). (E), (F), (G), (H) As opposed to MSTN inhibition, the rescue in long-term contractures with continuous bortezomib treatment in male mice at 8 and 12 weeks post-NBPI is associated with a decrease in β5 proteasome subunit activity, rather than β1 activity of denervated biceps muscles (n = 3 independent mice). In (B) and (C), elbow and shoulder contracture severity is calculated as the difference in passive elbow extension and shoulder rotation, respectively between the denervated (NBPI) side and the contralateral control side. In (D), sarcomere length on the NBPI side is normalized to the contralateral side to generate a sarcomere length ratio. Data are presented as mean ± SD. Statistical analyses: (B), (C), (D) 2-way ANOVA for sex and treatment with a Bonferroni correction for multiple comparisons at each time point, (E), (F), (G), (H) 3-way ANOVA for sex, treatment, and denervation (repeated measures between forelimbs) with a Bonferroni correction for multiple comparisons. **\***P<0.05, **\*\***P<0.01, ***P<0.001, **\*\*\*\***P<0.0001. This figure is generated from primary data reported in Goh et al., 2021.^13^

## Discussion

Secondary muscle contractures in pediatric neuromuscular disorders such as NBPI and cerebral palsy are major drivers of joint immobility, limb deformity, and physical dysfunction and disability.^4^ As existing treatments for contractures have focused on palliative mechanical solutions and do not address the underlying pathophysiology of contracture formation, the effectiveness of these strategies in restoring physical function is limited.^5–7^ Specifically, the paucity in our understanding of contracture pathophysiology impedes clinical efforts to prevent contractures. To overcome these limitations, we have previously identified the dysregulation of muscle proteostasis, as characterized by an increase in proteasome-mediated protein degradation, as a causative mechanism of impaired longitudinal muscle growth and contracture formation following NBPI.^12^ This discovery therefore highlights the possibility of pharmacologic strategies to prevent contractures through targeting of a biological mechanism. As our recent findings revealed a need for chronic pharmacologic proteasome inhibition with the potential for cumulative off-target toxicity,^13^ we must continue to identify safe and efficacious strategies for preventing pediatric muscle contractures by targeting muscle-specific regulators of proteostasis.

In our present study, we pharmacologically targeted MSTN, a prominent muscle-specific negative regulator of proteostasis, and identified several key findings regarding MSTN signaling in muscle growth and contracture pathophysiology. First, we discovered that pharmacologic MSTN inhibition with ACVR2B-Fc enhanced the size and protein content of normally innervated forelimb muscles in 1 month-old mice at P33, which is equivalent to the completion of neonatal muscle development in human infants.^35^ In comparison, the youngest age that prior studies on MSTN inhibition have reported gains in lean body mass was in 2 month-old mice.^36^ Our current finding therefore indicates a key regulatory role for MSTN signaling in governing muscle growth during the neonatal window, an often overlooked period in developmental muscle biology. This discovery not only reveals greater insights in skeletal muscle development, but also offers potential therapeutic strategies for overcoming low muscle mass arising from a host of genetic, metabolic, and hormonal disorders in the pediatric population.^37^ In addition, MSTN inhibition also protected against atrophy and sarcomere overstretch in neonatally denervated forelimb muscles, as well as prevented contractures in female but not male mice. These findings therefore establish a sex-specific role for MSTN as a driver of impaired muscle growth and contracture formation exclusively in female mice. Critically, this sex dimorphism lies in the pathophysiology of denervation-induced atrophy and contractures rather than drug pharmacokinetics or pharmacodynamics, given that MSTN inhibition promotes robust growth in normally innervated muscles of both female and male mice. Indeed, MSTN inhibition restored proteostasis only through the attenuation of 20S proteasomal subunit activity in denervated muscles of female mice, without associated changes in overall levels of protein synthesis or K48-linked polyubiquitin. This postulates an intriguing mechanism for MSTN-dependent impairment of longitudinal muscle growth, whereby MSTN putatively circumvents canonical signaling pathways of proteostasis to directly regulate the 20S proteasome after neonatal muscle denervation. Beyond offering potential translational opportunities for female NBPI patients, these seminal findings highlight the underappreciated role of sex as a biological variable in the pathophysiology and treatment of acquired neuromuscular disorders. Our results also establish that longitudinal muscle growth is governed through sex-divergent and noncanonical pathways, which offers valuable directions for future investigations to undertake.

Skeletal muscle exhibits high levels of sex dimorphisms. Female and male muscles typically display heterogeneity in body composition and protein turnover,^23,38^ regenerative and hypertrophic processes,^39,40^ metabolism and substrate utilization,^41,42^ drug pharmacodynamics,^43^ and even the development of disease-induced pathologies.^44^ While the role of sex in neonatal muscle denervation is largely unknown, we speculate that sex mediates divergent pathways that ultimately lead to contracture formation. Our current discovery of a sex-specific role for MSTN signaling in mediating contractures thus establishes a critical link between neonatal muscle denervation and impaired longitudinal muscle growth. In neonatally denervated female muscles, the improvement in functional muscle length following MSTN inhibition is associated with reduced proteasome subunit activity. This finding compelled us to thoroughly scrutinize our prior results on proteasome inhibition with bortezomib through the lens of sex differences.^13^ Here, we discovered that bortezomib treatment preferentially prevented elbow and shoulder contractures in male mice after the neonatal stage of muscle development and into skeletal maturity. Though the sex-associated phenotypic differences are more subtle in this context, they further illustrate sex-divergent pathways in the modulation of NBPI-induced contractures.

Intriguingly, the effects of MSTN inhibition on proteasome activity in female mice parallel our earlier results of proteasome inhibition with bortezomib.^13^ Specifically, we observed that a 30-45% attenuation in either β1 or β5 subunit activity at 4 weeks post-NBPI is associated with a 50-55% and a 45-60% reduction in elbow and shoulder contractures, respectively. Furthermore, this magnitude of reduction in either subunit activity is associated with an improvement of normalized sarcomere length by 14-15%. The absence of subunit specificity in preventing contractures thus points to a less prominent role for the different proteolytic sites of the 20S proteasome in modulating sarcomerogenesis and muscle length than previously proposed.^13^ Nevertheless, our collective findings indicate that different upstream pathways in contracture formation between sexes ultimately converge at the 20S proteasome, thereby validating the role of the proteasome as a key mediator of NBPI-induced contractures. Despite these insights, the precise mechanism(s) by which MSTN signaling modulates proteasome activity in denervated female muscles remains to be determined. While we observed altered Akt/mTOR and Smad2/3 signaling with NPBI, which are corroborated by studies from other groups using different mouse denervation models,^45–47^ MSTN inhibition failed to impact these canonical pathways in female mice. These findings thus suggest a non-canonical signaling mechanism through which MSTN regulates the 20S proteasome in female muscles after NBPI. Future studies are needed to elucidate this prospective alternative pathway. Further investigations into the molecular interactions between MSTN and proteasome activity are also critical for unraveling the complex regulatory machinery in longitudinal muscle growth, a poorly understood aspect of skeletal muscle biology.^48,49^ Additionally, it is unclear how proteostasis dysregulation and contractures are mediated in denervated male muscles. To this end, it is conceivable to postulate a role for sex hormones in contracture pathophysiology post-NBPI. An earlier study on juvenile frogs reported a loss of laryngeal muscle fibers only in male frogs following surgical denervation, whereas androgen treatment resulted in an additive effect on total fiber numbers only in innervated female laryngeal muscles.^50^ These results not only indicate a sexually dimorphic interaction between innervation and the endocrine system in regulating muscle fiber number, but they also suggest a protective effect of the female sex hormones against muscle fiber loss with denervation. Additional studies further verified the role of the female endocrine system in regulating muscle homeostasis. Ovariectomy in adult female rats prevented muscle mass recovery after hindlimb unloading,^51^ whereas genetic deletion of estrogen receptor β in muscle stem cells compromised muscle regeneration in juvenile female mice after local injections of barium chloride.^40^ Thus, to gain mechanistic insights on sexual dimorphisms in MSTN-mediated contracture formation, future studies must rigorously interrogate the contributions of sex in contracture pathophysiology by exploring the relationships between the different sex hormones and MSTN signaling in NBPI.

The finding of sex dimorphism in MSTN signaling itself is not without precedent. Targeted deletion of MSTN during skeletal maturity increased masseter mass and bite force in male mice only.^52^ In contrast, long-term MSTN deletion in skeletal muscles increased lean muscle mass only in aged female mice.^53^ These discrepancies might be attributable to sex-related differences of MSTN expression itself in skeletal muscles. Indeed, while men are prone to having higher expression of genes encoding ribosomal and mitochondrial proteins, women tend to have higher gene expression of the ACVR2B receptor.^54^ Moreover, there is an increased transcriptional and translational expression of processed MSTN in hindlimb muscles of adult female mice compared to their male counterparts.^55^ As the increased expression of MSTN and its receptor ligands facilitate higher MSTN activity in females, they conceivably predispose female muscles to more prominent responses from MSTN inhibition. In our current study, these intrinsic sex differences could potentially elucidate how pharmacologic MSTN inhibition confers greater neonatal growth in normally innervated female muscles and protects against atrophy in denervated muscles of female mice. These sex differences in MSTN and receptor gene expression may further account for the mixed outcomes of MSTN inhibition therapy for treating Duchenne Muscular Disease and other muscle disorders.^18,56^

Our discovery of sex dimorphisms in denervated muscle growth also calls into question prior conclusions on the role of MSTN signaling in denervation-induced atrophy from studies analyzing only male mice. In a juvenile mouse model of sciatic nerve resection, MacDonald et al. reported an inability of both rapamycin and ACVR2B-Fc treatment to prevent atrophy in male mice, and surmised that denervation atrophy is not Akt/mTOR or MSTN-dependent (2014).^46^ These findings are similar to our current observations of male mice in a neonatal mouse model of NBPI. In our present study, we extend MacDonald et al.’s earlier findings by revealing a novel role for MSTN in mediating atrophy solely in denervated muscles of female mice. Importantly, the sex dimorphisms in muscle atrophy are not limited to neonatal denervation, as sex differences in the pathophysiology of muscle atrophy have also been documented by an independent group using an adult mouse model of tenotomy.^57^ Overall, these latest discoveries underscore the need to account for sex as a biological variable in future studies investigating in different models of muscle atrophy.

Our study is not without limitations. Indeed, we observed that the ACVR2B-Fc decoy receptor attenuated growth of other non-skeletal muscle tissues, including the heart, spleen, and bone. Our observations are thus consistent with prior reports of off-target effects with pharmacologic MSTN inhibition.^18^ The off-target effects we observed in our study could have been minimized through genetic models to manipulate MSTN signaling in the skeletal muscle. While such genetic models could reveal additional insights in contracture formation, the off-target effects do not fully explain the sex-specificity of our outcomes. As MSTN plays a key role in cardiac energy homeostasis,^58^ the attenuation in heart size of male mice could impair cardiac function and lead to reduced muscle activity. We are unable to ascertain this possibility as we did not perform any metabolic measurements in the current study, though there were no observable differences in spontaneous cage activity with ACVR2B-Fc treatment. Of particular note is the effect of MSTN inhibition on bone growth, which could potentially confound our measurements of muscle parameters that are normalized to humerus length. However, the reduced bone length in male denervated limbs following MSTN inhibition would, if anything, cause an apparent increase in muscle size and/or rescue of contractures. The absence of such muscle findings despite this effect on bone growth reinforces the lack of a beneficial effect of MSTN inhibition on male denervated muscle, further underscoring the sex dimorphisms seen in muscle parameters. Finally, although we identified a link between MSTN inhibition and proteasome activity without perturbations in canonical MSTN signaling pathways, the molecular mechanisms by which MSTN could be regulating proteasome activity remain unclear. Potential noncanonical pathways could include p21-activated kinase 1 (PAK1) signaling, which has recently been shown to be permissive to gains in skeletal muscle mass with MSTN inhibition.^59^ Alternatively, MSTN might target contractures through the autophagy-lysosome pathway instead.^60^ Future studies are needed to fully elucidate mechanistic links and complete our understanding of the role of MSTN signaling in contracture pathophysiology.

## Conclusion

In conclusion, we established several critical insights in this current study that enhance our understanding of contracture pathophysiology and establish the framework for future investigations. First, the efficacy of MSTN inhibition at rescuing contractures, at least in female mice, reinforces the role of skeletal muscle biology in contracture formation and provides proof of concept for potential muscle-specific therapies for contracture prevention and treatment. Second, the sex-specific improvement in longitudinal muscle growth is associated with a corresponding reduction in proteasome activity, offering further evidence that the proteasome is a key regulator of muscle contractures. Third, the rescue in longitudinal growth without perturbations in the canonical signaling pathways with which MSTN regulates proteostasis suggests a potential noncanonical pathway governing muscle length. Lastly, the discovery of sex dimorphisms in our current study highlights the importance of sex in the pathophysiology of denervation atrophy, muscle growth, and pediatric muscle contractures. Moving forward, we strongly advocate the inclusion of sex as a biological variable in both mechanistic and translational studies on muscle atrophy and acquired neuromuscular diseases.

## Materials and Methods

### NBPI surgical mouse model

To assess the role of MSTN signaling in pediatric muscle contracture formation, we utilized our surgical mouse model of postganglionic NBPI where injury to the brachial plexus at postnatal day (P)5 induces contractures within 4 weeks after denervation.^12,13,61^ Briefly, unilateral, global (C5-T1), post-ganglionic NBPIs were surgically created in P5 wildtype female and male mice (Charles River; CD-1® IGS mouse, strain code 022) by extraforaminal nerve root excision under isoflurane anesthesia. Following the surgery, mice were returned to their mums and housed in standard cages with bio-huts on a 12-hour light/12-hour dark cycle, with nutrition and activity *ad libitum*. In order to ensure permanent deficits and to prevent potential confounding consequences of reinnervation, we validated deficits in motor function both postoperatively and prior to sacrifice. We excluded from the study mice that displayed preserved or recovered movement in the denervated limb. Based on this criteria, 1 female mouse was excluded from the study.

### MSTN inhibition

Beginning one day prior to surgery and continuing for four weeks after surgery, we pharmacologically inhibited the MSTN pathway by treating wildtype neonatal mice with a soluble decoy receptor fused to an Fc domain (ACVR2B-Fc).^19,20^ The binding of MSTN to the Fc domain inhibits MSTN from binding to Activin A, which blocks MSTN activity in the muscle, ultimately leading to increased muscle protein synthesis and robust muscle growth.^19^ The dosage frequency for the ACVR2B decoy receptor varies from 3,^20,21^ 5-6,^22^ and 7 days.^19,20^ Since this is the first known study in which this drug was tested in neonatal mice, we utilized the lowest dosage frequency in order to prevent potential toxicity.^19,20^

Hence, the soluble decoy receptor was administered weekly at a dose of 10 mg/kg via intraperitoneal injections, whereas Dulbecco’s phosphate-buffered saline (PBS) was used as a control in a separate litter of mice (**Figure 1A**). This regimen was performed twice to obtain 2 separate control groups and 2 corresponding drug-treated groups. All control and experimental groups in this study were randomized by litter, with all treatments administered at noon and in the respective cages. Milk spots in non-weaned mice and body weight at all ages were monitored daily for adequate nursing and signs of toxicity. Mice displaying an inability to nurse/self-feed post weaning were euthanized immediately.

### Assessment of contractures

Mice were euthanized at 4 weeks post-NBPI (P33) by CO_2_ asphyxiation, whereupon passive range of motion of the elbow and shoulder joints were assessed to determine contracture severity. Prior to sacrifice, mice were fasted for 4 hours and then administered 21.8 mg/kg puromycin (Sigma-Aldrich #P7255) intraperitoneally for 30 minutes, which gets incorporated into newly formed peptide chains.^25^ Immediately post-sacrifice, digital photography images of bilateral elbows and shoulders were captured at maximum passive extension and external rotation, respectively. Elbow flexion and shoulder internal rotation contractures were subsequently calculated in AxioVision (Zeiss).^12,13,61^ This previously validated method of assessing passive range of motion and contracture severity was performed with blinding to treatment groups. The forelimb images shown in **Figures 3A** and **3D** are representative of their respective samples, and have been processed to reflect comparable levels of sharpness, brightness, and contrast for illustrative purposes. However, no image manipulation was performed prior to measurements.

### Tissue collection and preparation

Following digital photography, we harvested bilateral biceps muscles (long head only), bilateral triceps muscles, hearts, and spleens. Biceps, hearts, and spleens were also weighed. Both biceps and triceps were then flash-frozen and stored at –80°C for subsequent analysis of protein dynamics and signaling pathways. The remaining forelimbs (with intact brachialis muscles) were positioned at 90° elbow flexion on cork, and imaged with digital radiographs for humerus length. Bilateral forelimbs were subsequently fixed in 10% formalin at 90° elbow flexion to avoid sarcomere relaxation with muscle removal. Following 48 hours of fixation, bilateral brachialis muscles were dissected, soaked in 25% Lugol solution (Sigma-Aldrich #32922) overnight, and imaged by micro computed tomography for assessment of whole muscle cross-sectional area and volume.^12,13,61^ Brachialis muscles were then recovered by soaking in PBS overnight at 4°C, digested in 15% sulfuric acid for 30 minutes to obtain muscle bundles, and imaged for sarcomere length.

### Ex vivo high-resolution studies

MicroCT was performed using a Siemens Inveon PET/SPECT/CT Scanner (Siemens Medical Solutions, Malvern, PA, USA) as previously described.^61^ The cone-beam CT parameters were as follows: 360° rotation, 1080 projections, 1300-ms exposure time, 1500-ms settle time, 80-kVp voltage, 500-µA current, and effective pixel size 17.67 µm. Briefly, acquisitions were reconstructed using a Feldkamp algorithm with mouse beam-hardening correction, slight noise reduction, and 3D matrix size 1024×1024×1536, using manufacturer-provided software. Protocol-specific Hounsfield unit (HU) calibration factor was applied.

### Muscle length

Although the functional length of a whole muscle is defined by its total number of sarcomeres in series, morphological constraints restrict its direct measurement.^12^ To overcome these limitations, we measured the average sarcomere length at 90° elbow flexion to determine the relative functional length of the brachialis muscles on control and denervated limbs. As previously described, elongated (overstretched) sarcomeres indicate fewer sarcomeres in series.^13^ This correlates with a shorter functional whole muscle length, because the fewer sarcomeres a muscle has in series, the more each sarcomere has to stretch to accommodate any given position.

Following overnight recovery in PBS at 4°C, the brachialis muscles were digested in 15% sulfuric acid for 30 minutes, and then dissected into muscle bundles for imaging with differential interference contrast (DIC) microscopy at 40x on a Nikon Ti-E SpectraX widefield microscope. Six images representing different muscle bundles of the same brachialis were acquired per muscle. Average sarcomere length of the brachialis was subsequently determined by measuring a series of 10 sarcomeres from each of the 6 representative DIC images with the AxioVision program as previously described,^12,13,61^ with blinding to treatment groups. Representative sarcomere images in **Figure 4A** have been cropped to identical sizes, and processed to reflect comparable levels of sharpness, brightness, and contrast for illustrative purposes. No image manipulation was performed prior to measurements.

### Humerus length and whole muscle size

To quantify humerus lengths from digital radiographs, AxioVision software was used to measure the distance between the proximal humerus physis to the distal articular surface.^12,13,61^ Whole muscle cross-sectional area and muscle volume of the brachialis were obtained by processing the MicroCT scans into digital imaging and communications in medicine (DICOM) images with Fiji programs (Segmentation Editor and 3D Viewer, respectively), and normalized to humerus length of the corresponding forelimb.^12,13,61^ All measurements were performed with blinding to treatment groups. For illustrative purposes, raw DICOM files were processed in IMARIS software (Bitplane, Zurich, Switzerland) to create whole-muscle images presented in **Figures 1B** and **2A**.

### Assessment of protein dynamics

To determine the effects of MSTN inhibition on muscle proteostasis, we assayed for protein synthesis by detection of the antibiotic puromycin using non-radioactive surface sensing of translation (SUnSET),^24,25^ and protein degradation by quantification of the amount of proteins marked for ubiquitination, with blinding to treatment groups.^12^ Bilateral biceps muscles were first bead homogenized (TissueLyser II; Qiagen) at a frequency of 30 Hz for 3 cycles of 2 minutes in lysis buffer [10 mM Tris (pH 7.4), 1 mM EDTA, 1 mM dithiothreitol, 0.5% Triton X-100, 2.1 mg/ml NaF] containing protease and phosphatase inhibitor cocktails (5 μl/ml; Sigma-Aldrich #P8340 and #P5726, respectively).^62^ Following centrifugation at 15,000 x g for 10 minutes at 4°C, the amount of protein in supernatants was quantified using Bradford protein assay. Biceps muscle homogenates were then heated at 65°C for 30 minutes, separated on 4-20% SDS-PAGE gels (30 µg of proteins), and transferred at 4°C to PVDF-FL membranes (Immobilon). Homogenates from adult mouse plantaris muscles that had been subjected to 7 days of mechanical overload were included in each gel as positive controls.^62^ Membranes were subsequently blocked in 5% BSA in TRIS-buffered saline (TBS)-Tween (pH 7.5), and incubated overnight at 4°C with an antibody against puromycin (1:1000; MilliporeSigma #MABE343), or K48-linkage-specific-polyubiquitin (1:1000, Cell Signaling #8081). GAPDH (1:5000; Cell Signaling #2118) served as a control for sample loading. Membranes were then washed and incubated with IRDye 800CW anti-mouse IgG2a (1:5000; LI-COR Biosciences) or DyLight^TM^ anti-rabbit (1:5000; Cell Signaling #5151) secondary antibodies. Following image detection on the Odyssey infrared detection system (LI-COR Biosciences), the relative abundance of puromycin incorporation and K48-linked protein levels were quantified using the Image Studio Lite program (LI-COR Biosciences), and normalized to corresponding GAPDH protein levels.^12,62^

To determine the effects of MSTN inhibition on proteasome activity, we assayed for proteasome subunit catalytic activity with blinding to treatment groups as previously described.^12,13^ Briefly, bilateral triceps muscles were homogenized in 20 mM of Tris-HCl (pH 7.2), 0.1 mM of EDTA, 1 mM of 2-mercaptoethanol, 5 mM of ATP, 20% of glycerol, and 10% (v/v) IGEPAL® CA-630 (Sigma-Aldrich #I8896).^63^ Following centrifugation described above, protein concentration was determined using the Pierce 660 nm protein assay kit (Thermo Scientific #22662). The caspase-like activity of the 20S proteasome β-1 catalytic subunit and chymotrypsin-like activity of the β-5 catalytic subunit were assayed with 10 μg total protein per muscle, through detection of the 7-amino-4-methylcoumarin (AMC) labeled fluorogenic peptide substrates Z-LLE-AMC (Boston Biochem #S-230) and LLVY-AMC (Chemicon #APT280), respectively. Assays were carried out in a white opaque polystyrene 96-well plate for 2 hours at 37°C, with endpoint fluorescence measured at 380 nm/460 nm on a SpectraMax M5 microplate reader (Molecular Devices). Relative fluorescence units were calculated per μg protein.

### Assessment of signaling pathways

To assess the effects of MSTN inhibition on downstream pathways, we assayed for Akt/mTOR and Smad2/3 signaling, with blinding to treatment groups. Briefly, triceps muscle homogenates were heated at 65°C for 30 minutes, separated on 10% SDS-PAGE gels (20 µg of proteins), and transferred at 4°C to PVDF-FL membranes (Immobilon).^62^ Membranes were subsequently blocked in 5% BSA/TBS-Tween, and incubated overnight at 4°C with an antibody against phosphorylated Akt (Ser473) (1:750; Cell Signaling #9271), total Akt (1:750; Cell Signaling #9272), phosphorylated Smad3 (Ser423, Ser425) (1:2000 Abcam #ab51451), or total Smad3 (1:2000; Abcam #ab28379). GAPDH (1:5000; Cell Signaling #2118) served as a control for sample loading. Membranes were then washed and incubated with a DyLight^TM^ anti-rabbit secondary antibody (1:5000; Cell Signaling #5151). Western blot signals were subsequently imaged and the relative abundance of phosphorylated and total protein levels of Akt and Smad3 were quantified as described above.

### Statistical analysis

All statistical tests were performed with Prism 8 software (GraphPad). For all continuous data sets that contained at least four independent samples, outliers were detected *a priori* by Grubbs’ test and excluded. To compare differences in values between two data sets, all data were subsequently tested for normality with the Shapiro–Wilk test. Normally distributed data were compared with unpaired two-tailed Student’s t-tests, whereas non-normally distributed data were compared using the Mann–Whitney U-tests. For data sets with two independent variables (sex and treatment), a 2-way ANOVA with Bonferroni correction for multiple comparisons was performed. For data sets with three independent variables (sex, treatment, denervation), a 3-way ANOVA with repeated measures between forelimbs and Bonferroni correction for multiple comparisons was performed. All data are presented as mean ± SD. The degree of significance between data sets is depicted as follows: *P < 0.05, **P < 0.01, ***P < 0.001, and ****P < 0.0001. *A priori* power analyses based on prior work determined that 6 mice per group were required to detect a 10° difference in contractures and a 0.2 µm difference in sarcomere lengths at 80% power between experimental conditions. A total of 32 mice (17 female-10 control, 7 experimental; and 15 male-8 control, 7 experimental) were used in this study.

### Ethical statement

This study was performed in strict accordance with recommendations in the Guide for the Care and Use of Laboratory Animals of the National Institutes of Health. All rodents were handled according to approved institutional animal care and use committee (IACUC) protocols (#2020-0067) of the Cincinnati Children’s Hospital Medical Center, and every effort was made to minimize suffering. This study also adhered to the ARRIVE 2.0 guidelines, and a checklist is provided with the manuscript.

### Material Availability Statement

All data generated or analyzed during this study are included in the manuscript and supporting files.

## Abbreviations

NBPI: neonatal brachial plexus injury
GDF8: growth differentiation factor-8
TGF-β: transforming growth factor beta
MSTN: myostatin
ACVR2B-Fc: soluble activin receptor type IIB
Akt: protein kinase B
mTOR: mammalian target of rapamycin
p70^S6K^: S6 kinase
Smad 2 & 3: mothers against decapentaplegic homolog 2 & 3
GAPDH: glyceraldehyde-3-phosphate dehydrogenase
DPBS: Dulbecco’s phosphate-buffered saline

## Acknowledgements

We thank the following entities within Cincinnati Children’s Hospital Medical Center: the Veterinary Services Core for surgical assistance, the Confocal Imaging Core for microscope assistance, and the Millay Laboratory for discussions and feedback. We also thank Sharon Wang from the Preclinical Imaging Core (University of Cincinnati College of Medicine) for MicroCT assistance. This work was supported by grants to RC from the National Institutes of Health (NIH) (R01HD098280-01), as well as funding from the Cincinnati Children’s Hospital Division of Orthopaedic Surgery and Junior Cooperative Society. The respective funding sources were not involved in the study design; in the collection, analysis, and interpretation of data; in the writing of the report; and in the decision to submit the paper for publication.

## Author Contributions

Q. Goh and R. Cornwall conceived and supervised the study; M.E. Emmert, Q. Goh, and R. Cornwall designed the research and experiments; M.E. Emmert, P. Aggarwal, K. Shay-Winkler, Q. Goh, and R. Cornwall performed research; S.J. Lee contributed new reagents; M.E. Emmert, P. Aggarwal, K. Shay-Winkler, S.J. Lee, Q. Goh, and R. Cornwall analyzed and/or interpreted data; M.E. Emmert and Q. Goh wrote the manuscript with assistance from all authors.

## Figure Legends

**Figure 1 – supplemental figure 1:**
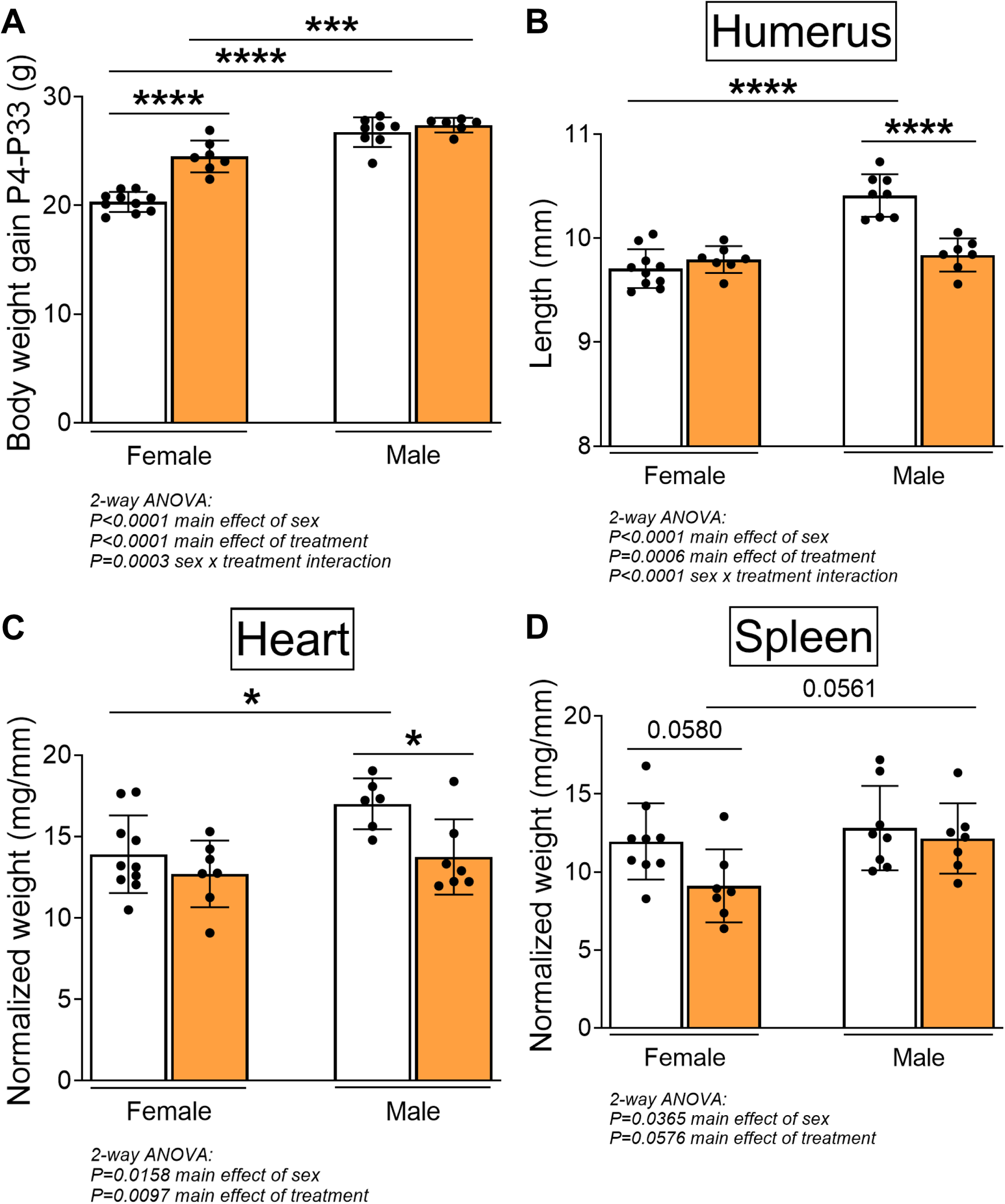
Off-target effects of pharmacologic MSTN inhibition. Treatment with ACVR2B-Fc (A) does not promote additional weight gains in male mice, but instead, (B) attenuates skeletal growth of non-denervated forelimbs and (C) reduces heart size. In contrast, ACVR2B-Fc treatment (D) reduces spleen size only in female mice. Data are presented as mean ± SD, n = 7-10 independent mice. Statistical analyses: (A), (B), (C), (D) 2-way ANOVA for sex and treatment with a Bonferroni correction for multiple comparisons. **\***P<0.05, **\*\*\***P<0.001, **\*\*\*\***P<0.0001.

**Figure 2 – supplemental figure 1:**
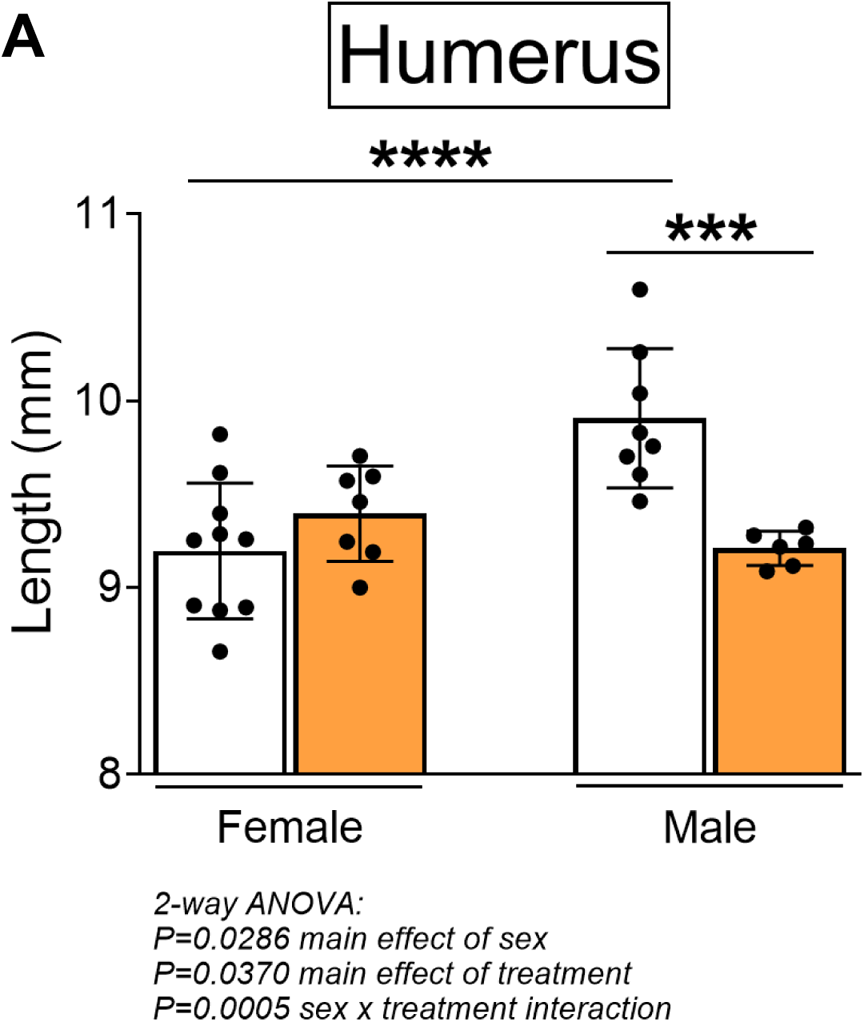
Effect of pharmacologic MSTN inhibition on skeletal growth in denervated forelimb. (A) Treatment with ACVR2B-Fc attenuates skeletal growth of denervated forelimbs only in male mice. Data are presented as mean ± SD, n = 7-10 independent mice. Statistical analyses: 2-way ANOVA for sex and treatment with a Bonferroni correction for multiple comparisons. **\*\*\***P<0.001, ****P<0.0001.

## References

1. Foad SL, Mehlman CT, Ying J. The epidemiology of neonatal brachial plexus palsy in the United States. J Bone Joint Surg Am. 2008;90(6):1258–64.

2. Govindan M, Burrows HL. Neonatal Brachial Plexus Injury. Pediatr Rev. 2019;40(9):494–6.

3. Pondaag W, Malessy MJ, van Dijk JG, Thomeer RT. Natural history of obstetric brachial plexus palsy: a systematic review. Dev Med Child Neurol. 2004;46(2):138–44.

4. Hale HB, Bae DS, Waters PM. Current concepts in the management of brachial plexus birth palsy. J Hand Surg Am. 2010;35(2):322–31.

5. Newman CJ, Morrison L, Lynch B, Hynes D. Outcome of subscapularis muscle release for shoulder contracture secondary to brachial plexus palsy at birth. J Pediatr Orthop. 2006;26(5):647–51.

6. Pedowitz DI, Gibson B, Williams GR, Kozin SH. Arthroscopic treatment of posterior glenohumeral joint subluxation resulting from brachial plexus birth palsy. J Shoulder Elbow Surg. 2007;16(1):6–13.

7. van der Sluijs JA, van Ouwerkerk WJ, de Gast A, Nollet F, Winters H, Wuisman PI. Treatment of internal rotation contracture of the shoulder in obstetric brachial plexus lesions by subscapular tendon lengthening and open reduction: early results and complications. J Pediatr Orthop B. 2004;13(3):218–24.

8. Nikolaou S, Peterson E, Kim A, Wylie C, Cornwall R. Impaired growth of denervated muscle contributes to contracture formation following neonatal brachial plexus injury. J Bone Joint Surg Am. 2011;93(5):461–70.

9. Weekley H, Nikolaou S, Hu L, Eismann E, Wylie C, Cornwall R. The effects of denervation, reinnervation, and muscle imbalance on functional muscle length and elbow flexion contracture following neonatal brachial plexus injury. J Orthop Res. 2012;30(8):1335–42.

10. Nikolaou S, Liangjun H, Tuttle LJ, Weekley H, Christopher W, Lieber RL, et al. Contribution of denervated muscle to contractures after neonatal brachial plexus injury: not just muscle fibrosis. Muscle Nerve. 2014;49(3):398–404.

11. Nikolaou S, Hu L, Cornwall R. Afferent Innervation, Muscle Spindles, and Contractures Following Neonatal Brachial Plexus Injury in a Mouse Model. J Hand Surg Am. 2015;40(10):2007–16.

12. Nikolaou S, Cramer AA, Hu L, Goh Q, Millay DP, Cornwall R. Proteasome inhibition preserves longitudinal growth of denervated muscle and prevents neonatal neuromuscular contractures. JCI Insight. 2019;4(23).

13. Goh Q, Nikolaou S, Shay-Winkler K, Emmert ME, Cornwall R. Timing of proteasome inhibition as a pharmacologic strategy for prevention of muscle contractures in neonatal brachial plexus injury. FASEB J. 2021;35(2):e21214.

14. Lee SJ. Regulation of muscle mass by myostatin. Annu Rev Cell Dev Biol. 2004;20:61–86.

15. Sartori R, Milan G, Patron M, Mammucari C, Blaauw B, Abraham R, et al. Smad2 and 3 transcription factors control muscle mass in adulthood. Am J Physiol Cell Physiol. 2009;296(6):C1248–57.

16. Trendelenburg AU, Meyer A, Rohner D, Boyle J, Hatakeyama S, Glass DJ. Myostatin reduces Akt/TORC1/p70S6K signaling, inhibiting myoblast differentiation and myotube size. Am J Physiol Cell Physiol. 2009;296(6):C1258–70.

17. Lee SJ. Myostatin: Regulation, function, and therapeutic applications. Muscle. 2012;2:1077–84.

18. Lee SJ. Targeting the myostatin signaling pathway to treat muscle loss and metabolic dysfunction. J Clin Invest. 2021;131(9).

19. Lee SJ, Reed LA, Davies MV, Girgenrath S, Goad ME, Tomkinson KN, et al. Regulation of muscle growth by multiple ligands signaling through activin type II receptors. Proc Natl Acad Sci U S A. 2005;102(50):18117–22.

20. Lee SJ, Huynh TV, Lee YS, Sebald SM, Wilcox-Adelman SA, Iwamori N, et al. Role of satellite cells versus myofibers in muscle hypertrophy induced by inhibition of the myostatin/activin signaling pathway. Proc Natl Acad Sci U S A. 2012;109(35):E2353–60.

21. Goh Q, Song T, Petrany MJ, Cramer AA, Sun C, Sadayappan S, et al. Myonuclear accretion is a determinant of exercise-induced remodeling in skeletal muscle. Elife. 2019;8.

22. Lee SJ, Lehar A, Meir JU, Koch C, Morgan A, Warren LE, et al. Targeting myostatin/activin A protects against skeletal muscle and bone loss during spaceflight. Proc Natl Acad Sci U S A. 2020;117(38):23942–51.

23. Griffin GE, Goldspink G. The increase in skeletal muscle mass in male and female mice. Anat Rec. 1973;177(3):465–9.

24. Goodman CA, Mabrey DM, Frey JW, Miu MH, Schmidt EK, Pierre P, et al. Novel insights into the regulation of skeletal muscle protein synthesis as revealed by a new nonradioactive in vivo technique. FASEB J. 2011;25(3):1028–39.

25. Schmidt EK, Clavarino G, Ceppi M, Pierre P. SUnSET, a nonradioactive method to monitor protein synthesis. Nat Methods. 2009;6(4):275–7.

26. Bodine SC, Stitt TN, Gonzalez M, Kline WO, Stover GL, Bauerlein R, et al. Akt/mTOR pathway is a crucial regulator of skeletal muscle hypertrophy and can prevent muscle atrophy in vivo. Nat Cell Biol. 2001;3(11):1014–9.

27. Rommel C, Bodine SC, Clarke BA, Rossman R, Nunez L, Stitt TN, et al. Mediation of IGF-1-induced skeletal myotube hypertrophy by PI(3)K/Akt/mTOR and PI(3)K/Akt/GSK3 pathways. Nat Cell Biol. 2001;3(11):1009–13.

28. Egerman MA, Glass DJ. Signaling pathways controlling skeletal muscle mass. Crit Rev Biochem Mol Biol. 2014;49(1):59–68.

29. Sartori R, Romanello V, Sandri M. Mechanisms of muscle atrophy and hypertrophy: implications in health and disease. Nat Commun. 2021;12(1):330.

30. McFarlane C, Plummer E, Thomas M, Hennebry A, Ashby M, Ling N, et al. Myostatin induces cachexia by activating the ubiquitin proteolytic system through an NF-kappaB-independent, FoxO1-dependent mechanism. J Cell Physiol. 2006;209(2):501–14.

31. Lokireddy S, Mouly V, Butler-Browne G, Gluckman PD, Sharma M, Kambadur R, et al. Myostatin promotes the wasting of human myoblast cultures through promoting ubiquitin-proteasome pathway-mediated loss of sarcomeric proteins. Am J Physiol Cell Physiol. 2011;301(6):C1316–24.

32. Yau R, Rape M. The increasing complexity of the ubiquitin code. Nat Cell Biol. 2016;18(6):579–86.

33. Chen JL, Walton KL, Hagg A, Colgan TD, Johnson K, Qian H, et al. Specific targeting of TGF-beta family ligands demonstrates distinct roles in the regulation of muscle mass in health and disease. Proc Natl Acad Sci U S A. 2017;114(26):E5266–E75.

34. Goodman CA, McNally RM, Hoffmann FM, Hornberger TA. Smad3 induces atrogin-1, inhibits mTOR and protein synthesis, and promotes muscle atrophy in vivo. Mol Endocrinol. 2013;27(11):1946–57.

35. Dutta S, Sengupta P. Men and mice: Relating their ages. Life Sci. 2016;152:244–8.

36. McPherron AC, Lee SJ. Suppression of body fat accumulation in myostatin-deficient mice. J Clin Invest. 2002;109(5):595–601.

37. Orsso CE, Tibaes JRB, Oliveira CLP, Rubin DA, Field CJ, Heymsfield SB, et al. Low muscle mass and strength in pediatrics patients: Why should we care? Clin Nutr. 2019;38(5):2002–15.

38. Smith GI, Mittendorfer B. Sexual dimorphism in skeletal muscle protein turnover. J Appl Physiol (1985). 2016;120(6):674–82.

39. Kitajima Y, Ono Y. Estrogens maintain skeletal muscle and satellite cell functions. J Endocrinol. 2016;229(3):267–75.

40. Seko D, Fujita R, Kitajima Y, Nakamura K, Imai Y, Ono Y. Estrogen Receptor beta Controls Muscle Growth and Regeneration in Young Female Mice. Stem Cell Reports. 2020;15(3):577–86.

41. Rosa-Caldwell ME, Greene NP. Muscle metabolism and atrophy: let’s talk about sex. Biol Sex Differ. 2019;10(1):43.

42. Maher AC, Fu MH, Isfort RJ, Varbanov AR, Qu XA, Tarnopolsky MA. Sex differences in global mRNA content of human skeletal muscle. PLoS One. 2009;4(7):e6335.

43. Salamone IM, Quattrocelli M, Barefield DY, Page PG, Tahtah I, Hadhazy M, et al. Intermittent glucocorticoid treatment enhances skeletal muscle performance through sexually dimorphic mechanisms. J Clin Invest. 2022.

44. De Jonghe B, Sharshar T, Lefaucheur JP, Authier FJ, Durand-Zaleski I, Boussarsar M, et al. Paresis acquired in the intensive care unit: a prospective multicenter study. JAMA. 2002;288(22):2859–67.

45. Castets P, Rion N, Theodore M, Falcetta D, Lin S, Reischl M, et al. mTORC1 and PKB/Akt control the muscle response to denervation by regulating autophagy and HDAC4. Nat Commun. 2019;10(1):3187.

46. MacDonald EM, Andres-Mateos E, Mejias R, Simmers JL, Mi R, Park JS, et al. Denervation atrophy is independent from Akt and mTOR activation and is not rescued by myostatin inhibition. Dis Model Mech. 2014;7(4):471–81.

47. Tando T, Hirayama A, Furukawa M, Sato Y, Kobayashi T, Funayama A, et al. Smad2/3 Proteins Are Required for Immobilization-induced Skeletal Muscle Atrophy. J Biol Chem. 2016;291(23):12184–94.

48. Jorgenson KW, Hornberger TA. The Overlooked Role of Fiber Length in Mechanical Load-Induced Growth of Skeletal Muscle. Exerc Sport Sci Rev. 2019;47(4):258–9.

49. Jorgenson KW, Phillips SM, Hornberger TA. Identifying the Structural Adaptations that Drive the Mechanical Load-Induced Growth of Skeletal Muscle: A Scoping Review. Cells. 2020;9(7).

50. Tobias ML, Marin ML, Kelley DB. The roles of sex, innervation, and androgen in laryngeal muscle of Xenopus laevis. J Neurosci. 1993;13(1):324–33.

51. Sitnick M, Foley AM, Brown M, Spangenburg EE. Ovariectomy prevents the recovery of atrophied gastrocnemius skeletal muscle mass. J Appl Physiol (1985). 2006;100(1):286–93.

52. Williams SH, Lozier NR, Montuelle SJ, de Lacalle S. Effect of Postnatal Myostatin Inhibition on Bite Mechanics in Mice. PLoS One. 2015;10(8):e0134854.

53. Tavoian D, Arnold WD, Mort SC, de Lacalle S. Sex differences in body composition but not neuromuscular function following long-term, doxycycline-induced reduction in circulating levels of myostatin in mice. PLoS One. 2019;14(11):e0225283.

54. Welle S, Tawil R, Thornton CA. Sex-related differences in gene expression in human skeletal muscle. PLoS One. 2008;3(1):e1385.

55. McMahon CD, Popovic L, Jeanplong F, Oldham JM, Kirk SP, Osepchook CC, et al. Sexual dimorphism is associated with decreased expression of processed myostatin in males. Am J Physiol Endocrinol Metab. 2003;284(2):E377–81.

56. Rybalka E, Timpani CA, Debruin DA, Bagaric RM, Campelj DG, Hayes A. The Failed Clinical Story of Myostatin Inhibitors against Duchenne Muscular Dystrophy: Exploring the Biology behind the Battle. Cells. 2020;9(12).

57. Meyer GA, Thomopoulos S, Shen KC. Tenotomy-induced muscle atrophy is sex-specific and independent of NFκβ. Manuscript in co-submission.

58. Biesemann N, Mendler L, Wietelmann A, Hermann S, Schafers M, Kruger M, et al. Myostatin regulates energy homeostasis in the heart and prevents heart failure. Circ Res. 2014;115(2):296–310.

59. Barbe C, Loumaye A, Lause P, Ritvos O, Thissen JP. p21-Activated Kinase 1 Is Permissive for the Skeletal Muscle Hypertrophy Induced by Myostatin Inhibition. Front Physiol. 2021;12:677746.

60. Wang DT, Yang YJ, Huang RH, Zhang ZH, Lin X. Myostatin Activates the Ubiquitin-Proteasome and Autophagy-Lysosome Systems Contributing to Muscle Wasting in Chronic Kidney Disease. Oxid Med Cell Longev. 2015;2015:684965.

61. Ho BL, Goh Q, Nikolaou S, Hu L, Shay-Winkler K, Cornwall R. NRG/ErbB signaling regulates neonatal muscle growth but not neuromuscular contractures in neonatal brachial plexus injury. FEBS Lett. 2021;595(5):655–66.

62. Goh Q, Millay DP. Requirement of myomaker-mediated stem cell fusion for skeletal muscle hypertrophy. Elife. 2017;6.

63. Penna F, Bonetto A, Aversa Z, Minero VG, Rossi Fanelli F, Costelli P, et al. Effect of the specific proteasome inhibitor bortezomib on cancer-related muscle wasting. J Cachexia Sarcopenia Muscle. 2016;7(3):345–54.

